# Agency and responsibility over virtual movements controlled through different paradigms of brain–computer interface

**DOI:** 10.1101/735548

**Authors:** Birgit Nierula, Bernhard Spanlang, Matteo Martini, Mireia Borrell, Vadim V. Nikulin, Maria V. Sanchez-Vives

## Abstract

Agency is the attribution of an action to the self and is a prerequisite for experiencing responsibility over its consequences. Here we investigated agency and responsibility by studying the control of movements of an embodied avatar, via brain computer interface (BCI) technology, in immersive virtual reality. After induction of virtual body ownership by visuomotor correlations, healthy participants performed a motor task with their virtual body. We compared the passive observation of the subject’s ‘own’ virtual arm performing the task with (1) the control of the movement through activation of sensorimotor areas (motor imagery) and (2) the control of the movement through activation of visual areas (steady-state visually evoked potentials). The latter two conditions were carried out using a brain–computer interface (BCI) and both shared the intention and the resulting action. We found that BCI-control of movements engenders the sense of agency, which is strongest for sensorimotor areas activation. Furthermore, increased activity of sensorimotor areas, as measured using EEG, correlates with levels of agency and responsibility. We discuss the implications of these results for the neural bases of agency, but also in the context of novel therapies involving BCI and the ethics of neurotechnology.

**Key points summary:**

- We induced embodiment of a virtual body and its movements were controlled by two different BCI paradigms – one based on signals from sensorimotor versus one from visual cortical areas.
- BCI-control of movements engenders agency, but not equally for all paradigms.
- Cortical sensorimotor activation correlates with agency and responsibility.
- This has significant implications for neurological rehabilitation and neuroethics.

## Introduction

The feeling of generating actions that can influence the course of events in the outside world is a central feature of human experience and is important for our perception of the self (Haggard, 2017). The attribution of actions to ourselves is referred to as “agency” (Gallagher, 2000), which gives us a feeling of control and results in a sense of responsibility over an action’s outcome.

Through brain-computer interfaces (BCI) it is feasible to control the actions of surrogate bodies, prostheses, robots, or other effectors. Such BCI systems are based on a closed feedback loop, in which neural activity related to an intention is transformed in real-time into a command to a device that then gives feedback to the participant (Birbaumer, 2006; Hochberg et al., 2012; Pfurtscheller & Neuper, 2001). Different types of BCI are available, some of which rely on changes in the sensorimotor mu-rhythm induced through motor imagery. For such motor-imagery-related BCI it has been shown that discrepancies between the neural activity and the resulting visual feedback can result in a decreased sense of agency (Evans, Gale, Schurger, & Blanke, 2015; Marchesotti et al., 2017). However, whether and how agency can be modulated by the usage of other BCI systems has not been investigated.

Agency is closely linked to body ownership—the feeling that “this body is my body”—and although agency and body ownership can be dissociated (Imaizumi & Asai, 2015; Kalckert & Ehrsson, 2012; Tsakiris, Longo, & Haggard, 2010), agency has been shown to significantly correlate with body ownership (Kalckert & Ehrsson, 2012).

Both body ownership and agency are aspects of self-consciousness, which might not be a unique elemental percept or qualia but rather a cluster of subjective experiences, feelings, and attitudes (Synofzik, Vosgerau, & Newen, 2008; Tsakiris et al., 2010). The neural underpinnings of agency have been extensively investigated (for an overview see David (2012) and Haggard (2017)) and might lie in the connectivity between several brain areas, rather than in any single brain structure. A possible candidate is the connectivity between areas involved in action initiation and areas involved in monitoring of perceptual events, which have been proposed as neural correlates (Haggard, 2017). For events caused by an external agent (external agency), neuroimaging studies report activation of a complex brain network including the dorsomedial prefrontal cortex, precuneus, temporoparietal junction and pre-supplementary motor area (pre-SMA). However, for agency-related activity the insula seems to be the most consistently reported area (Haggard, 2017; Sperduti, Delaveau, Fossati, & Nadel, 2011). On the other hand, interference with activity in the pre-SMA by means of transcranial stimulation has been shown to alter agency (Cavazzana, Penolazzi, Begliomini, & Bisiacchi, 2015; Moore, Ruge, Wenke, Rothwell, & Haggard, 2010), pointing to an involvement of motor areas in the development of the sense of agency. The role of motor areas in the sense of agency is theoretically supported by a computational model of agency, the comparator model (Frith, Blakemore, & Wolpert, 2000), which is based on a model for sensorimotor integration (Wolpert, Ghahramani, & Jordan, 1995) and describes agency as the result of a matching between predicted and actual sensory feedback of a planned motor action (Blakemore, Wolpert, & Frith, 2000; Frith et al., 2000). Further, brain regions related to BCI control not related to the comparator model have been identified in the basal ganglia, the anterior cingulate cortex and the left superior frontal gyrus (Marchesotti et al., 2017).

Under certain conditions, we can perceive agency over an agent’s action without performing that action with our own body, which is referred to as “illusory agency”. For example, one can perceive agency over another person’s hand movements when seeing that hand at the position where one would expect one’s own hand to be and when there is a prior command to carry out the later seen actions (Wegner, Sparrow, & Winerman, 2004). Experiments on agency have also been conducted in immersive virtual environments, where a virtual body can be felt as one’s own after inducing adequate sensorimotor correlations (Sanchez-Vives, Spanlang, Frisoli, Bergamasco, & Slater, 2010; Slater, Perez-Marcos, Ehrsson, & Sanchez-Vives, 2008; Slater, Pérez-Marcos, Ehrsson, & Sanchez-Vives, 2009). Impressively, during this “virtual embodiment” experience, people may attribute actions of the virtual body to themselves even though they are not performing those actions with their real body (Banakou & Slater, 2014; Kokkinara, Kilteni, Blom, & Slater, 2016).

Drawing upon these premises, our objective was to investigate whether and how the sense of agency over a virtual limb movement controlled through a BCI was generated. In order to generate a movement through BCI there is first an intention. Next, the brain signal is used by the BCI to initiate the movement, however different signals (or paradigms) can be used. Here we questioned: do agency levels differ between different BCI methods? How relevant is the cortical area that the BCI uses for the initiation of the movement? And, if agency over a BCI-controlled action is induced, is there also a feeling of responsibility with respect to the results of that action?

To answer these questions we designed and carried out an experiment to measure cortical activation patterns, sense of agency, and the sense of responsibility with respect to movements of an “embodied” virtual body in an immersive virtual environment (Slater et al., 2008; Slater, Spanlang, Sanchez-Vives, & Blanke, 2010) and compared three different paradigms to initiate the movements of a virtual arm: motor imagery, activating sensorimotor areas, (Pfurtscheller, Brunner, Schlögl, & Lopes da Silva, 2006; Pfurtscheller & Neuper, 1997); visually evoked potentials, activating visual cortical areas (Middendorf, McMillan, Calhoun, & Jones, 2000; Vialatte, Maurice, Dauwels, & Cichocki, 2010); and passive, non-triggered movements.

## Methods

### Participants

Sixty-two healthy, right-handed females with normal or corrected-to-normal vision, no neurological or psychological disorders, and no medication intake that could influence their perception were screened for the experiment. After the screening procedure, a sample of 40 participants completed the experiment. All participants were novices to BCI. To maintain a similar level of accuracy for BCI conditions, we used the inverse of the binomial cumulative distribution for finding the threshold for non-random (at *P*=0.001) classification of epochs. This threshold was in our case 30/40=0.75 (i.e. 75%), leaving a remaining sample of 29 participants (mean age=21.5 years, *SD=*2.6; laterality quotient (Oldfield, 1971): mean*=*71.0, *SD=*23.4). All participants were naïve to the research question, gave written informed consent before starting the experiment, and received €5–20 for their participation. The experiment was approved by the local ethics committee (Comité Ético de Investigación Clínica de la Corporación Sanitaria, Hospital Clínic de Barcelona) and is in accordance with the Declaration of Helsinki.

### Virtual environment and tracking system

The virtual environment was programmed and controlled using the Unity game engine (Unity Technologies, www.unity3d.com) and displayed through a HMD (Oculus Rift Development Kit 2, www.oculus.com) with a nominal field-of-view of 100°, a resolution of 960 × 1080 pixels per eye, and a 75 Hz frame rate. The virtual bodies were taken from the Rocketbox library (Rocketbox Studios GmbH, www.rocketbox.de). Further details about the laboratory setups to create and measure virtual embodiment illusions can be found in Spanlang et al. (2014). The virtual room was a custom-made replica of a virtual reality laboratory. During the embodiment phase, movements of the right hand were tracked with the right arm setup of Perception Neuron (Noitom, Beijing, China), a full-body tracking system.

### Electroencephalographic recordings

Electrographic data were captured with 64 active Ag/AgCl ring electrodes (g.LADYbird, g.tec, Schiedlberg, Austria), amplified with g.HIamp (g.tec) and recorded with Matlab R2013a Simulink software (The MathWorks, Inc., Natrick, USA) on a separate computer. During the data acquisition, signals were band-pass filtered between 0.5 and 250 Hz, notch filtered at 50 Hz, and digitized at a rate of 512 Hz. Fifty-nine EEG electrodes were mounted on a g.Gamma cap (g.tec) and had standard positions in accordance with the 10-percent electrode system (Chatrian, Lettich, & Nelson, 1985). The ground electrode was located at AFz. The two electrooculographic (EOG) electrodes were placed next to the outer canthus and below the right eye. EOG and EEG electrodes were referenced to the right ear. The two electromyographic (EMG) electrodes had a bipolar montage and were located over the distal section and belly of the anterior deltoid muscle. Data were analyzed offline using Matlab and the Berlin Brain-Computer Interface toolbox (https://github.com/bbci/bbci_public) (Benjamin Blankertz et al., 2016) and online using Matlab Simulink. To compensate for network latencies between the two computers, we synchronized them by using the network time protocol and time-stamped triggers with the times of the Unity and the Matlab computer. Triggers were then adapted in the analysis so that the time stamps would match.

### SMR-based BCI

The sensorimotor rhythm (SMR)-based BCI used the Common Spatial Pattern (CSP) approach (B Blankertz, Tomioka, Lemm, Kawanabe, & Muller, 2008; Fukunaga, 1990; Graimann & Pfurtscheller, 2006; Koles, Lind, & Soong, 1995; Müller-Gerking, Pfurtscheller, & Flyvbjerg, 1999) to extract event-related desynchronization (ERD) over sensorimotor areas from 27 channels (see Fig. S1) and was provided by g.tec (“Common Spatial Patterns 2-class BCI”, V2.14.01). This is an established approach for distinguishing two classes of motor imagery, which were in our case imagination of a right arm pressing a button (similar to the movement they saw in the virtual environment) and a left foot movement (extension and flexion of the ankle joint). The foot movement served to create the classifier.

### Motor imagery training

Before the actual experiment, participants undertook motor imagery training consisting of two parts, both outside virtual reality. In the first part we taught participants how to imagine the movement in order to produce the greatest changes in the SMR (this took up to 25 min), and in the second part participants did up to four training sessions with the BCI, which followed the training procedure recommended by g.tec (Guger, Edlinger, Harkam, Niedermayer, & Pfurtscheller, 2003). In the majority of the participants one training session was done directly before the motor imagery condition.

### SSVEP-based BCI

The SSVEP-based BCI extracted the signal from eight occipital channels (see Fig. S1), was provided by g.tec (“SSVEP BCI”, V2.14.01), and modified for its use in a virtual environment. SSVEP stimuli were two buttons in the virtual environment; one blinked at a frequency of 7.5 Hz and the other at 9.4 Hz. An extra, dummy frequency (20 Hz) was also used during the SSVEP training phase to be able to form a better classifier.

### SSVEP training

The SSVEP-training was performed while participants wore the HMD and was adapted from Guger et al., (2012). Each frequency required five training trials and took about 4 min. Only the frequency of interest was displayed during the training phase; during the experiment, both frequencies (7.5 and 9.5 Hz) were displayed at the same time.

### Experimental procedure

All instructions were pre-recorded and given to participants through headphones. Preparation of EEG electrodes took about 40 min. Before starting the experiment, participants did the training for the SMR- and SSVEP-based BCIs. The classifiers of both training programs were calculated using gBSanalyze (Version 5.16.00, g.tec).

### Executed Movement

Executed Movement was recorded in 18 out of the 29 participants in order to capture the brain activity for each participant while moving their real right arm. Through headphones participants heard a beep every 8 s with a jitter of ± 1 s. They were instructed to move their right hand and press a button in front of them (within 2 s) whenever they heard a beep, performing a movement that resembled the virtual movement.

### Experimental conditions

Participants were asked to sit comfortably on a chair with their arms lying on a table in front of them keeping a distance of 55 cm between their middle fingers (Fig. 1A, Fig. S2). Each experimental condition started with an embodiment phase followed by the action phase. In both phases participants saw a virtual body from a first person perspective in the same position and collocated with their real body.

**Fig. 1.**
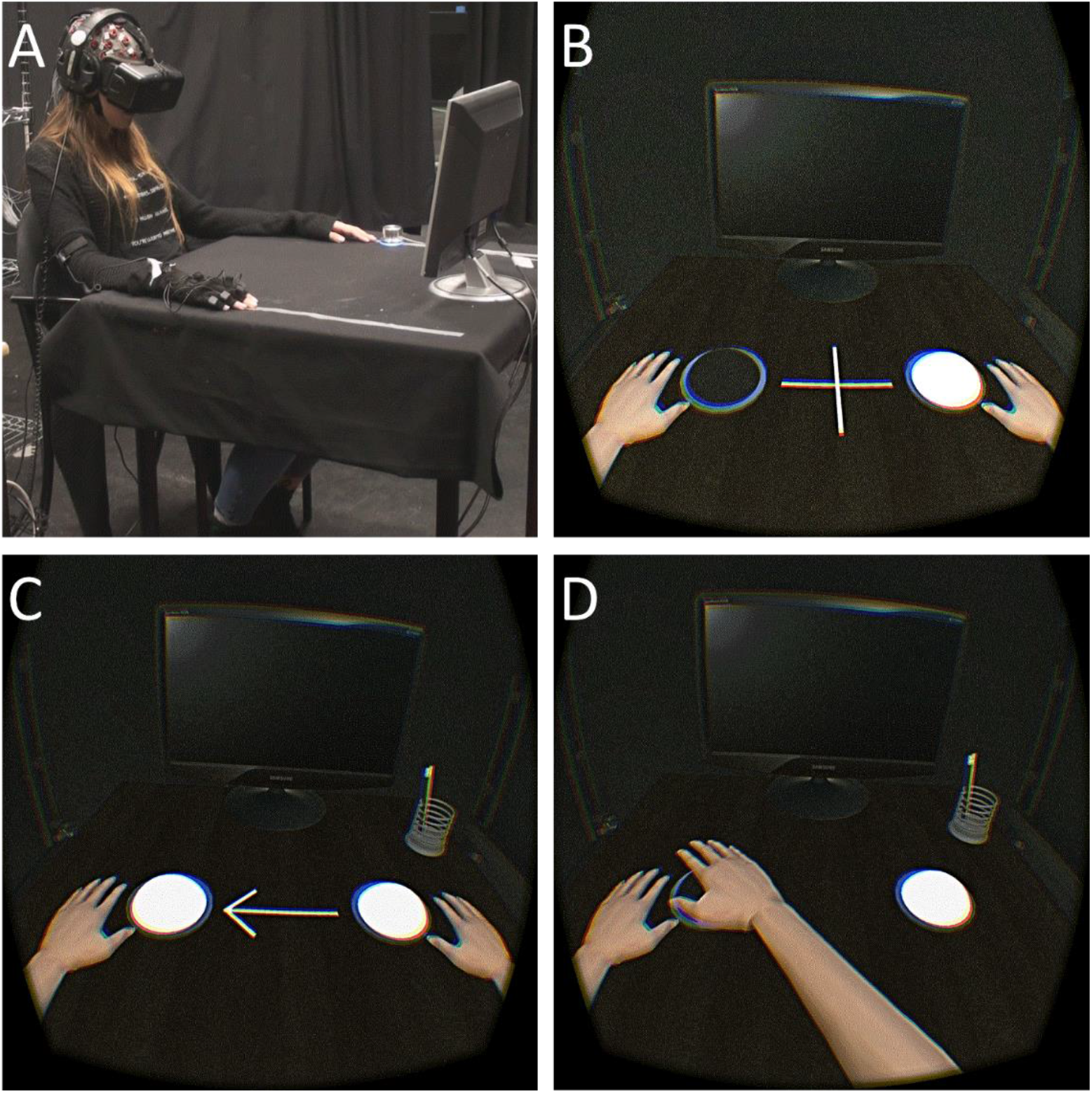
Experimental setup. (A) Participants sat with both hands resting on a table and saw through a head mounted display (HMD) a virtual body co-located with their real body, over which body ownership was induced in an initial embodiment phase. (B) During the action phase, in which participants were instructed not to move their arms, each trial started with a fixation cross, (C) followed by an arrow indicating one of two buttons to indicate the task, which was to bring the hand towards to pointed to-button. (D) In two conditions participants controlled the movement of the virtual arm through a BCI and in a third control condition they simply observed the virtual arm move.

#### Embodiment phase

When visualizing the virtual environment, participants were instructed to look around and describe what they saw, thereby inducing a feeling of presence in the virtual environment. They were further instructed to look down at the virtual body. Next, they were instructed to move their right hand and fingers for 1 min. These movements were tracked and mapped onto the movements of the virtual body, so that participants saw the right virtual arm moving in the same way. This procedure was carried out to induce a sense of body ownership and agency over the virtual body regardless of the condition. Directly after this, the questionnaire items IControlledArm and MyBody were presented on the virtual screen in randomized order (see Table 1).

**Table 1.**
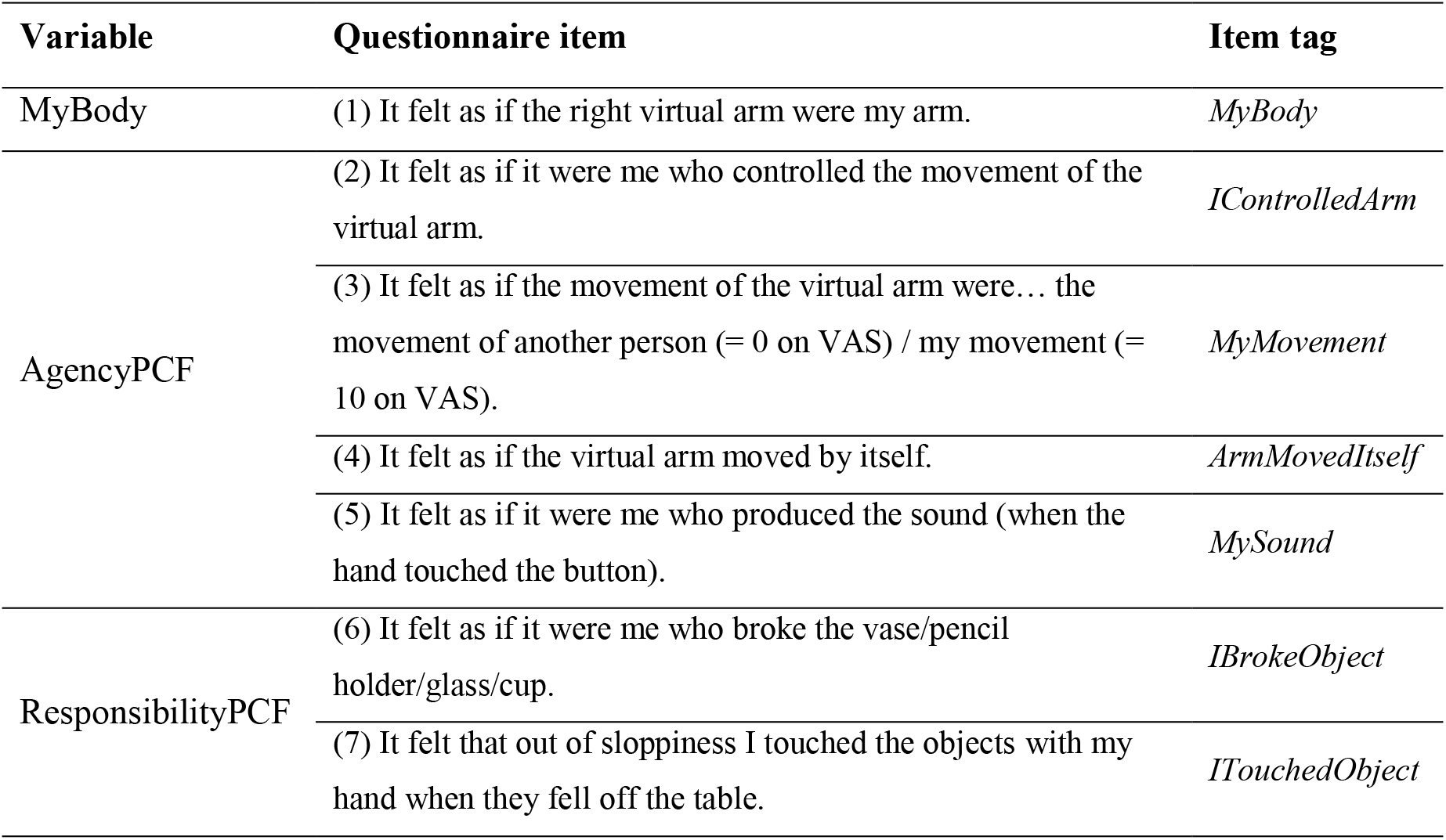
Questionnaire items answered by participants before, during, or after each experimental condition (in Spanish in the original). Responses were given on a visual analogue scale (VAS) ranging from 0 (“strongly disagree”) to 10 (“strongly agree”), except for item 3. Note that reversed wording was used for item 4 and therefore the inverse of the original scale was used for further analysis.

| Variable | Questionnaire item | Item tag |
| --- | --- | --- |
| MyBody | (1) It felt as if the right virtual arm were my arm. | *MyBody* |
| AgencyPCF | (2) It felt as if it were me who controlled the movement of the virtual arm. | *IControlledArm* |
|  | (3) It felt as if the movement of the virtual arm were... the movement of another person (= 0 on VAS) / my movement (= 10 on VAS). | *MyMovement* |
|  | (4) It felt as if the virtual arm moved by itself. | *ArmMovedItself* |
|  | (5) It felt as if it were me who produced the sound (when the hand touched the button). | *MySound* |
| ResponsibilityPCF | (6) It felt as if it were me who broke the vase/pencil holder/glass/cup. | *IBrokeObject* |
|  | (7) It felt that out of sloppiness I touched the objects with my hand when they fell off the table. | *ITouchedObject* |

#### Action phase

Directly after the embodiment phase, participants were instructed to keep still for the rest of the experimental condition and to only move the left hand when giving responses to the questionnaire items. The virtual environment was the same as during the embodiment phase, except that now they saw two buttons on the table in front of them (Fig. S3), each blinking at one of the SSVEP frequencies.

All participants went through the following three conditions: the MotorImagery condition, the SSVEP condition, and the Observe condition, each consisting of 40 trials. Each trial started with a fixation cross displayed between the two buttons (Fig. 1B), followed 2 s later by an arrow indicating the target button for that particular trial (Fig.1C). In the following 6 s participants had to either imagine the arm movement (MotorImagery condition), focus on the indicated button (SSVEP condition), or simply observe what was happening (Observe condition). During MotorImagery and SSVEP conditions the previously trained classifiers detected whether participants imagined the arm movement (MotorImagery) or to which button they looked (SSVEP); that is, if the frequency of the indicated button or if sensorimotor activity in arm areas were detected, the virtual arm would move toward the target button (Fig. 1D); if not, participants would hear an error sound to indicate a missed trial and the arm would not move. Missed trials were repeated one additional time. In the Observe condition, the arm moved in each trial to the target button. Fig. S3 displays the timeline of one trial. Every 10 trials participants responded to the IControlledArm item (Table 1). During each experimental condition the virtual arm accidentally threw an object off the table resulting in a breaking sound (Fig. S4). This happened four times throughout each condition at a random trial once during each block of 10 trials (but not directly before or after they were answering the IControlledArm item). The objects were a pencil stand, a vase, a coffee cup, and a glass, and they appeared in random order. Directly after the virtual arm broke an object, participants answered the IBrokeObject item (Table 1). One of the four trials was a catch trial which was included to test our items’ internal validity.

### Questionnaires

Table 1 displays all questionnaire items. Some items were displayed inside the virtual reality environment (MyBody, IControlledArm, IBrokeObject) and others outside on a computer screen (MyBody, MySound, ITouchedObject, MyMovement, ArmMovedItself). In the horizontal visual analogue scale for item 3, 0 corresponded to “the movement of another person” and 10 to “my movement” (see Table 1). Note that we used reverse wording when designing question 4, therefore its scale was reversed for the analysis.

Questionnaire items inside the virtual environment were displayed on a virtual screen. Participants gave their responses by turning a button (Power Mate, Griffin Technologies, Nashville, USA) located next to their left hand. VAS ratings, indicated by a little red bar, increased when the button was turned right and decreased when it was turned left. Participants confirmed their rating by pressing the button. In order to avoid interfering with the sense of body ownership, while rating the questionnaire items and therefore moving the left hand, the view of the virtual forearms was blocked by a virtual board.

Directly after each condition participants answered five questionnaire items outside the virtual environment, which were displayed on a laptop screen. In this case, participants answered the items using the keyboard. Questionnaire items were displayed in random order (see Table 1). Using a principal component factor analysis which resulted in one variable, the items IBrokeObject and ITouchedObject were merged to the variable ResponsibilityPCF, and the items IControlledArm, MyMovement, and ArmMovedItself were merged to the variable AgencyPCF. Initially we included also MySound, but excluded it later, because the resulting variable AgencyPCF explained more variance without this item (81% of variance *versus* 75% when including MySound). The continuous ratings of the MyBody item were down-scaled into ordinal ratings ranging from 1 to 10.

### Processing of EEG data

#### Artifact identification

Very noisy EEG channels were visually identified and removed from further analysis (this affected on average one to two channels in five participants). During data collection, one electrode (F5) broke and we continued without it (this affected 12 participants). For artifact detection the signal was filtered between 0.5 Hz and 40 Hz. Eye blink artifacts were identified and projected from EEG data with Independent Component Analysis (ICA) performed on concatenated epoched data (FastICA (Hyvärinen & Oja, 2000)). Epochs were generated around the onset of the arrow and consisted of 1000 ms of pre-stimulus and 6000 ms of post-stimulus interval. The epochs were visually inspected for the presence of muscular and mechanical artifacts, indicated by the variance of the maximal activity and by the Mahalanobis distance (Nikulin, Hohlefeld, Jacobs, & Curio, 2008). Within different subjects we identified between 0% and 8% of epochs.

### Executed Movement

Artifacts in the Executed Movement data were visually identified in epochs ranging from −1000 to 3000 ms around the tone indicating the start of the movement and those epochs containing them were removed at a later step in the analysis (0–25% of epochs).

#### Amplitude modulation of spontaneous alpha oscillations

We were interested in alpha oscillations because they have the largest signal-to-noise ratio of all spontaneous oscillations and therefore allow a reliable estimation of ERD (Nierula, Hohlefeld, Curio, & Nikulin, 2013; Nikouline et al., 2000; Pfurtscheller & Lopes da Silva, 1999), which is important for obtaining clear spatial patterns. Moreover, alpha oscillations have been previously shown to be a reliable indicator of engagement of cortical areas in different experimental conditions including sensory and motor tasks (Berger, 1929; Gastaut & Bert, 1954; Pfurtscheller & Aranibar, 1977; Pfurtscheller & Lopes da Silva, 1999).

#### Identifying individual sensorimotor alpha range

The individual sensorimotor alpha peak frequencies were identified in the pre-stimulus spectrum of Laplace transformed channel C3 (Graimann & Pfurtscheller, 2006; Hjorth, 1975). The signal of all conditions and the executed movement was then filtered in a range of ± 2 Hz around the individual peak frequency.

#### Identifying individual ERD peak latencies in executed movement data

The filtered and Laplace transformed signal was Hilbert transformed in order to obtain its amplitude envelope (Clochon, Fontbonne, Lebrun, & Etevenon, 1996; Graimann & Pfurtscheller, 2006; Rosenblum & Kurths, 1998). The signal was next cut into epochs from −1000 to 3000 ms around the tone and epochs containing artifacts were removed from further analysis. ERD% was calculated using the following equation: ERD% (*t*)= (AMP (*t*) – PRE) / PRE · 100, where AMP (amplitude) refers to the activity at each time point *t* of the averaged epochs and PRE is the mean amplitude in the pre-stimulus interval (from −500 to 0 ms). Time points with lowest ERD% were identified between 0 and 1500 ms in Laplace transformed channel C3 (*n=*18, latency: mean*=*808 ms, *SD=*228 ms).

#### Extracting CSP in Executed Movement data

The filtered, epoched, and cleaned executed movement signal was used for CSP analysis. No ICA was applied because prior application of another spatial filtering could lead to possible deterioration in the performance of CSP (Blankertz et al., 2008; Nierula et al., 2013). For CSP, pre-stimulus epochs, filtered in the individual alpha range, were merged from executed movement and all three experimental conditions, and the mean was subtracted from single epochs in the post-stimulus interval. The CSP algorithm is commonly used to separate two classes of data by determining the spatial filters W that maximize the variance of one class while simultaneously minimizing the variance of another class. In our data, one class contained the data of the pre-stimulus interval (from −500 to 0 ms) and the other class contained the data of the post-stimulus interval (±250 ms around the ERD peak previously identified in Laplace transformed C3 channel). In the case of bandpass-filtered EEG-signals, variance is equivalent to power in a given frequency range (Benjamin Blankertz, Dornhege, Krauledat, Müller, & Curio, 2005; Dornhege, Blankertz, & Curio, 2003; Nierula et al., 2013; Nikulin et al., 2008), which means that CSP can be also used to optimize ERD (Lemm, Müller, & Curio, 2009). CSP has been successfully used to classify single EEG epochs during motor imagery (Benjamin Blankertz et al., 2005; Dornhege et al., 2003; Koles et al., 1995; Nikulin et al., 2008). The inverse of the filter matrix W is the CSP and it contains components/patterns that were sorted by the size of their eigenvalue from high to low. Within the first four patterns we selected the pattern with the strongest eigenvalue that represented activity in sensorimotor areas. Patterns were validated by splitting all epochs into two sets. Only patterns that appeared in both sets were considered for further analysis.

#### Source reconstruction of CSP patterns

Source reconstruction of CSP patterns was performed using Brainstorm (Tadel, Baillet, Mosher, Pantazis, & Leahy, 2011). The forward model was generated with OpenMEEG using the symmetric Boundary Element Method (Gramfort, Papadopoulo, Olivi, & Clerc, 2010; Kybic et al., 2005) on the cortical surface of a Montreal Neurological Institute (MNI) brain template (non-linear MNI-ICBM152 atlas (Fonov et al., 2011)) with a resolution of 1 mm. Cortical sources were then estimated using the Tikhonov-regularized minimum-norm (Baillet, Mosher, & Leahy, 2001) with a Tikhonov parameter of *λ=*10% of maximum singular value of the lead field, and mapped to a distributed source model consisting of 15 000 current dipoles. Sources of CSP patterns were visually inspected for their origin in sensorimotor areas. Fig. 3A displays the inverse solutions averaged over all 18 participants, which was strongest at coordinates *X*=−52.5, *Y*=−7.5 and *Z*=52.9 in the MNI space. Mean sources were located in the precentral gyrus (Fig. 3A).

#### ERD

The extracted spatial filter *W* from the CSP pattern used for source analysis was also used to project the data *X* from the three conditions (MotorImagery, SSVEP, and Observe) using the following formula: *Z*=*WX*. To obtain ERD%, we extracted the amplitude envelope of the analytic signal with the Hilbert transform and then calculated ERD% (Nierula et al., 2013) using the formula above. The minimum value of ERD% during the preparation phase was later used for statistical comparisons.

#### EMG and EOG

The EMG recordings were high-pass filtered at 10 Hz and EOG recordings were low-pass filtered at 50 Hz. Both filtered EMG and EOG recordings were then segmented into epochs from −2000 to 8000 ms with respect to the arrow indicating one of the two buttons. The root-mean square values were calculated during the motor preparation phase (from 0 to 6000 ms) to obtain measures of motor activation (MA) in the right anterior part of the deltoid muscle and eye movement (EM).

MA and EM served as covariates in order to control for experiences of agency triggered by attributing eye or muscle movements to the virtual movement. Therefore, before fitting a statistical model we checked whether MA or EM could explain some of the variance.

### Statistics

All statistical tests were performed using Stata 13 (StataCorp LP, College Station, TX). Residuals were tested for normality when necessary using the Shapiro–Wilk test. Violations of the sphericity assumption were tested with Mauchly’s sphericity test. IBrokeObject and ITouchedObject were merged to the variable ResponsibilityPCF, and the items IControlledArm, MyMovement, and ArmMovedItself were merged to the variable AgencyPCF using a principal component factor analysis. The resulting interval scaled variables were analyzed with a multilevel mixed-effects regression (the “mixed” function in Stata) with “condition” as fixed-effects and “individual subject” as random effects. For MyBody, AgencyPCF, and ResponsibilityPCF likelihood ratio tests were used to identify if the variables MA or EM should be added as covariates to the model. For post hoc analyses we used the Scheffé post hoc criterion for significance. ERD% was analyzed using repeated measures ANOVA with factor “condition” and post hoc comparisons were performed using the Scheffé criterion. Because of its ordinal nature, MyBody reports were analyzed with a multilevel mixed-effects ordered logistic regression (the “meologit” function in Stata) with “condition” as fixed effects and “individual subject” as random effects. Relationships between variables were assessed with mixed-effects regression (the “mixed” function in Stata).

## Results

### Agency ratings

Agency has been measured in the literature as the sense of control over an action (Synofzik et al., 2008), as the sense of having caused/produced an outcome (Bednark & Franz, 2014; Kumar & Srinivasan, 2014), and as the sense that an action is one’s own and not another person’s (Weiss, Tsakiris, Haggard, & Schütz-Bosbach, 2014). In order to have a single measure for agency, we performed a principal component factor analysis on the questionnaire items indicating agency (IControlledArm, MyMovement, and ArmMovedItself; see Table 1), which resulted in a single variable accounting for 81% of variance. We refer to this variable as AgencyPCF.

A mixed-effects regression revealed an effect of condition on AgencyPCF (*z=*6.64, *P<*0.001), indicating that participants experienced higher levels of AgencyPCF over the virtual movement during the Motor Imagery than during the Observe condition (*Scheffé z=*6.64, *P*<0.001) and during the SSVEP conditions (*Scheffé z=*2.53, *P=*0.041), and participants experienced higher levels of AgencyPCF during the SSVEP than during the Observe condition (*Scheffé z=*4.12, *P<*0.001). Our data show that activating motor areas to induce the movement of the virtual arm elicited a higher sense of agency over the action than activating visual areas, and merely observing the virtual arm move elicited the lowest sense of agency (see Fig. 2A).

**Fig. 2.**
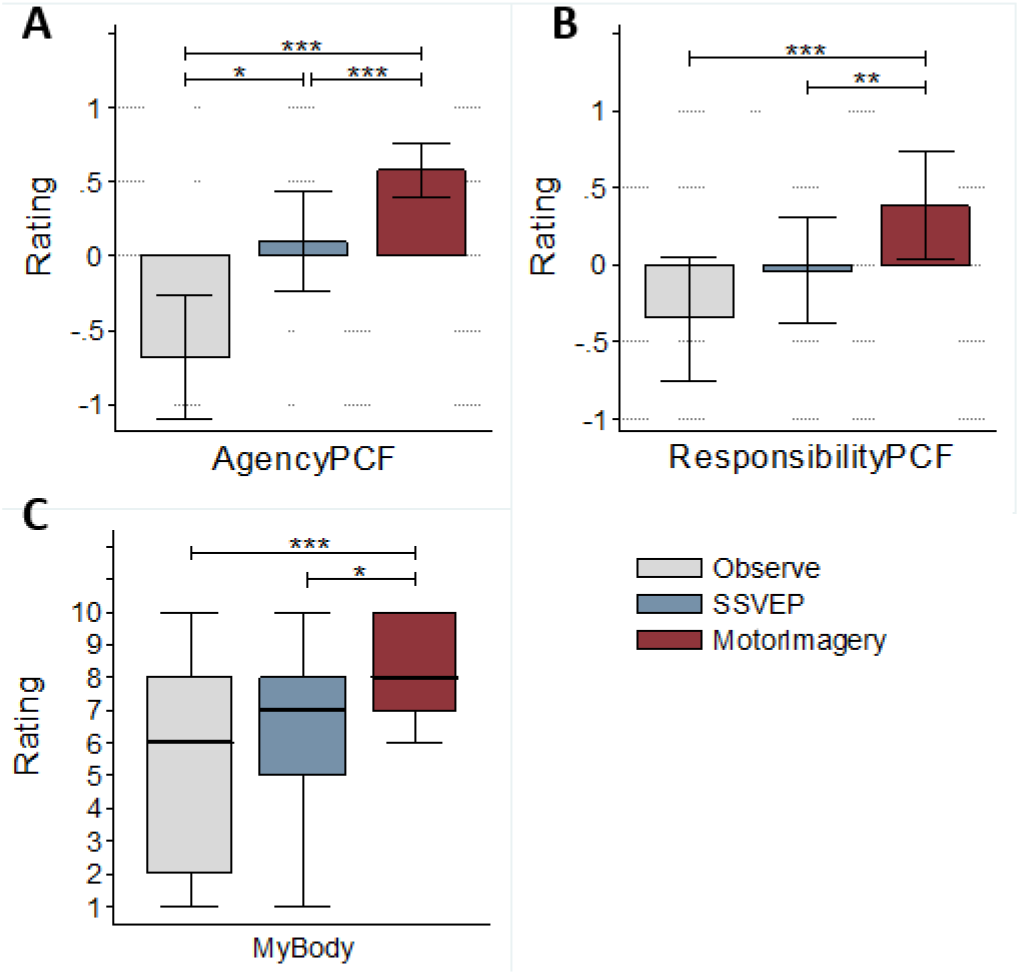
Agency, responsibility and body ownership ratings in the three experimental conditions (passive observation, SSVEP-BCI and motor imagery-BCI). AgencyPCF (A), ResponsibilityPCF (B), and MyBody (C) ratings. The Boxplots include all participants (n=29) who completed all conditions. The definition of these variables can be found in Results and Methods. See Table 1 for the questionnaire items associated with the variables. *: *P* ≤ 0.05, **: *P* ≤ 0.01, ***: *P* ≤ 0.001.

**Fig. 3.**
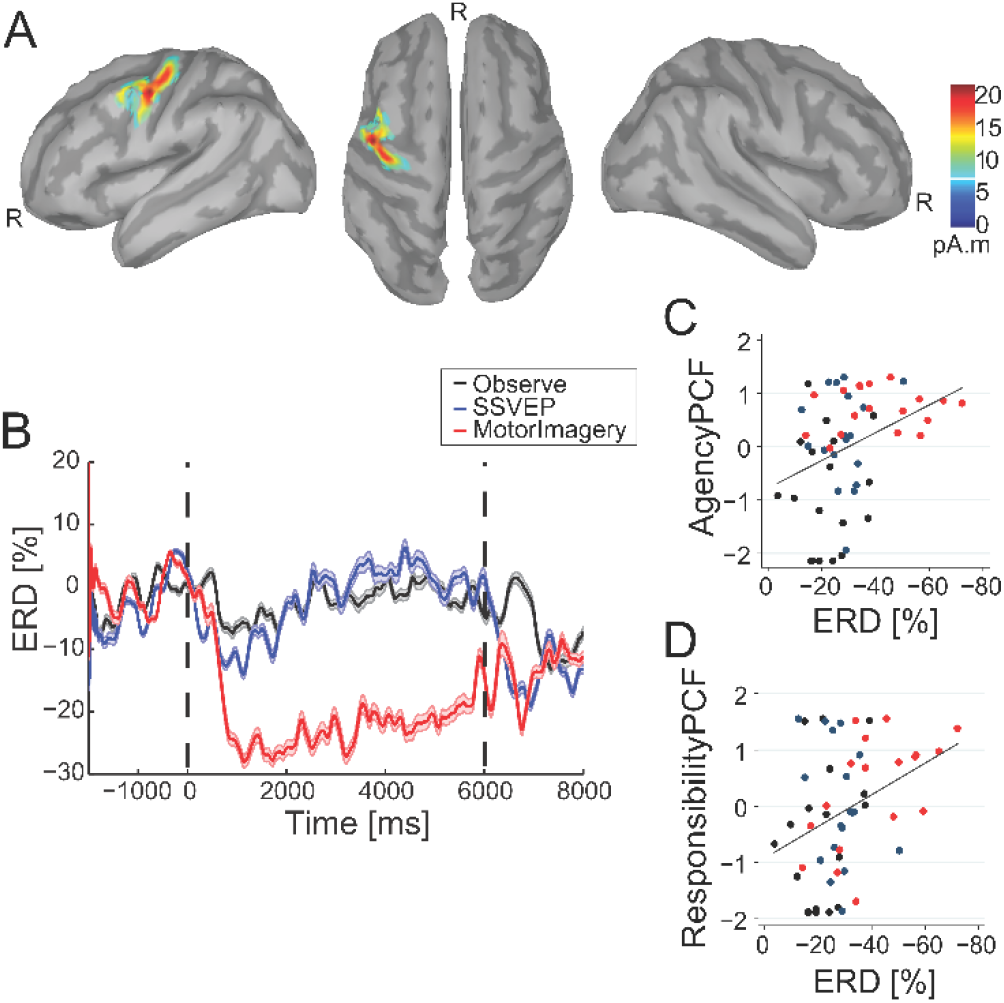
Brain activation during BCI-controlled and passive movements of the virtual arm and its relation to agency and responsibility. (A) Average sources of Executed Movement from 18 participants. The active area during this task coincides with the precentral gyrus (Brodmann’s area 6). The images display left hemisphere, top view with both hemispheres, and right hemisphere from left to right. R=rostral. (B) Event-related desynchronization (ERD) of the alpha rhythm as a marker of sensorimotor activation in the three conditions MotorImagery, SSVEP, and Observe. The signals, recorded during the three conditions, were projected through spatial filters, obtained by applying CSP on executed movement data, and transformed to ERD%. Zero (0) indicates the display of the arrow, after which participants started to observe, look at the blinking light or imagining the movement (depending on the condition). After 6000 ms the virtual arm started moving—in the Observe condition in every trial and in the BCI conditions when the action was classified accordingly. The plotted signal is averaged over n=18 participants. Only correctly classified trials were taken into account. (C) ERD% during Observe (in black), SSVEP (in blue), and MotorImagery (in red) plotted against AgencyPCF, (D) and ResponsibilityPCF. Note that the x-axis is inverted. More negative ERD% values indicate higher sensorimotor cortex activity. Lines indicate the linear fit of the questionnaire item on ERD%.

### Responsibility ratings

Agency is closely related to the feeling of responsibility because it tells us when we are responsible for the results of action and when we are not (Haggard & Tsakiris, 2009). To measure the levels of experienced responsibility, we introduced for each condition four random trials in which the virtual arm “accidentally” threw an object off the table and broke it (Supplementary Information, video). We asked participants directly after this incident how strongly they felt responsible for breaking the object (item IBrokeObject). One of these four trials was a catch trial, which we used to test if our measure was working. In this trial the object fell off the table although the virtual hand did not touch it. This trial was not part of the IBrokeObject measure (see Methods for more details). Moreover, once the condition (Motor Imagery, SSVEP or Observe) was over, participants were also asked whether they considered they had touched the object out of sloppiness (item ITouchedObject). We merged these two responsibility-related questionnaire items to the variable ResponsibilityPCF using a principal component factor analysis. A mixed-effects regression revealed an effect of condition on ResponsibilityPCF (*z=*4.66, *P<*0.001), wherein participants gave higher ratings during MotorImagery than during Observe (*Scheffé z=*4.66, *P*<0.001) or during SSVEP (*Scheffé z=*3.34, *P=*0.004), whereas there was no difference between the SSVEP and the Observe conditions (*Scheffé z=*1.18, not significant, n.s.). Our results show therefore that activating motor areas to induce the virtual arm movement led to a significantly higher sense of responsibility than using visual areas or merely observing the movement (Fig. 2B).

### Body ownership ratings

Body ownership and agency are closely related. For example, only when feeling high levels of body ownership over a surrogate body can people experience illusions of agency over the actions of that body (Banakou & Slater, 2014; Kokkinara et al., 2016) or even perceive action errors as their own (Padrao, Gonzalez-Franco, Sanchez-Vives, Slater, & Rodriguez-Fornells, 2016). To capture the participant’s experienced level of body ownership we asked them to rate it at the end of each condition (see Table 1 questionnaire item MyBody). A mixed-effects ordered logistic regression revealed an effect of condition on MyBody (*z=*3.40, *P=*0.001; Fig. 2), showing that ratings were highest during Motor Imagery compared to Observe (*Scheffé z=*3.40, *P=*0.001) and SSVEP (*Scheffé z=*2.22, *P=*0.026); during the Observe and SSVEP conditions ratings did not differ (*Scheffé z=*1.37, n.s). Our data show that participants had highest levels of body ownership during Motor Imagery, while during SSVEP and Observe conditions they reported body ownership at similarly intermediate levels (Fig. 2C).

### Activity in sensorimotor areas

Agency has been proposed to result from the match between predicted and actual sensory feedback of a planned motor action, a process that involves sensorimotor areas (Frith et al., 2000; Giummarra, Gibson, Georgiou-Karistianis, & Bradshaw, 2008). To test the different levels of sensorimotor activity induced by the experimental conditions we first recorded EEG activity in 18 of the 29 included participants while they pressed a button with their right arm (a movement that replicates the virtual one). Applying common spatial pattern (CSP) analysis to the executed movement data, we obtained spatial filters of sensorimotor activity, which we used to project the EEG signal of the three conditions and then calculate the event-related desynchronization (ERD) of the somatosensory alpha rhythm in 18 participants (Fig. 3B displays ERD% of the projected signal). We compared minimum ERD% in the different conditions during the preparation phase, which corresponds to a time window of 6 s before participants saw the virtual movement and in which they either imagined the movement, focused at the blinking light, or simply observed. A repeated measures analysis of variance (ANOVA) with fixed-effect condition revealed that there was an effect of condition on ERD% (*F (2,34)=*12.70, *P<*0.001), wherein ERD% was highest during MotorImagery compared to SSVEP (*Scheffé t=*–3.29, *P=*0.009) and Observe (*Scheffé t=*–4.95, *P<*0.001). The SSVEP and Observe conditions did not differ (*Scheffé t=*– 1.66, n.s.) (Table 1). Our data show that our experimental paradigm triggered processing in the sensorimotor cortex during the preparation phase and that corresponding levels of neural activity were highest during MotorImagery, and low during SSVEP and Observe conditions.

In order to relate sensorimotor activity to the sense of agency, as has been proposed by computational models of agency (Blakemore et al., 2000; Frith et al., 2000) as well as models that take a cognitive component of agency into account (Synofzik et al., 2008), we investigated the relationship between ERD% and agency-related questionnaire reports. We found that greater sensorimotor activity, which is reflected in negative ERD%, was associated with a higher sense of agency and responsibility. Specifically, a mixed-effects regression revealed a negative relationship between ERD% and both AgencyPCF (Fig. 3C; *z=*–3.02, *P=*0.003) and ResponsibilityPCF (Fig. 3D; *z=*–4.56, *P<*0.001). There was no significant relationship between ERD% and MyBody (*z=*–2.22, *P=*0.026). We then examined whether these relationships were also present within conditions. A mixed-effects regression revealed a negative relationship between ERD% and ResponsibilityPCF (*z=*– 2.84, *P=*0.004) in the MotorImagery condition, but not in the two other conditions (Observe: *z=*–1.27, n.s.; SSVEP: *z=*1.33, n.s.), indicating that when participants intentionally activated sensorimotor areas, the strength of sensorimotor activity was related to their feeling of responsibility; that is, the stronger the activity the higher the feeling of responsibility. There was no relationship between ERD% and AgencyPCF in any of the three conditions (Observe: *z=*–0.10, n.s.; SSVEP: *z=*–0.25, n.s.; Motor Imagery: *z=*–0.50, n.s.) and no relationship between ERD% and MyBody (Observe: *z=*–0.70, n.s.; SSVEP: *z=*1.97, *P*=0.048; MotorImagery: *z=*–1.13, n.s.). Our data therefore demonstrates that the stronger the sensorimotor areas' activation, the higher the subjective experience of agency and responsibility.

### Debriefing

Finally, in order to get an overall idea of how much control participants felt over the virtual movement during the different conditions, we asked them to give an overall rating at the end of the experiment. For this rating, participants were asked to order the conditions from strongest to weakest according to their experienced sense of control over the virtual arm movement (IControlledArm). Out of 29 participants, 22 reported strongest feelings of control during Motor Imagery, followed by SSVEP, and during the Observe condition the lowest feeling of control over the virtual arm movement was reported (Fig. S5).

## Discussion

With the present work we have demonstrated that BCI-controlled movements engender agency, being higher during this conditions than during the observation of a passive movement of the ‘own’ body. Furthermore, the highest agency is induced by BCI based on the activation of sensorimotor areas (motor imagery paradigms). Interestingly, BCI based on activation of visual areas (SSVEP paradigms), engendered elevated levels of agency compared to movement observation, but not a higher sense of responsibility. Our analysis also revealed that the higher the activation of sensorimotor areas, the higher the reported levels of agency and responsibility. The implications of these findings should be considered at different levels: (1) on the neural basis of agency and responsibility for actions, (2) on executive aspects of which BCI paradigms should be used for different purposes, and (3) on the ethical implications of controlling actions by thinking and the subjective responsibility over the consequences that this may imply.

### Higher agency but not responsibility levels for intended movements

Voluntary actions can be described by an intention–action–outcome chain (Roskies, 2010). In this chain, intentions are linked to actions by an action selection process, which is the basis for the motor plan. In addition to the selected action, the motor plan also computes its expected sensory feedback (Frith et al., 2000), and a comparison between intended and expected sensory feedback has been proposed to produce a sense of initiation (de Vignemont & Fourneret, 2004). The action selection process occurs after the formation of the intention but before its execution through an action. In an experimental setup priming can be used to bias the action selection process by making some actions easier to access than others. Participants report reduced feelings of agency when such breaks between the intention–action link are present (Valérian Chambon, Sidarus, & Haggard, 2014; Valerian Chambon, Wenke, Fleming, Prinz, & Haggard, 2013; Wenke, Fleming, & Haggard, 2010). Actions are also linked to their outcome through a comparator mechanism (Frith et al., 2000), which has been proposed to create a sense of one’s own movement and contribute, together with the sense of initiation (de Vignemont & Fourneret, 2004) and other cognitive processes (Synofzik et al., 2008), to the sense of agency. One difference between inducing the movement of a virtual limb through BCI and merely observing the movement is that in the latter condition no intention is generated, while to operate a BCI it is necessary to generate intention. Therefore, BCI-control should lead to a greater sense of agency than mere observation. Our data confirmed this prediction for perceived levels of agency but not for responsibility. Participants perceived higher levels of agency when an intention was generated before seeing the virtual movement, such as in the two BCI conditions, compared to when there was no intention; that is, when participants observed the movement (see Fig. 2). On the other hand, participants’ feelings of responsibility over the consequences of a given action, such as the virtual arm accidentally breaking an object in the virtual scenario, were higher only when they induced the virtual movement by the activation of sensorimotor areas. Contrarily, when the movement was induced by visually-evoked potentials the feeling of responsibility was similar to when participants simply observed the movement without being able to control it. One possible interpretation is that the intention to perform a ask, *per se*, is not a determining factor for the feeling of responsibility to arise.

### Importance of body ownership

Illusory agency is a phenomenon that can also give clues about the neural basis of agency. When experiencing illusory agency, one attributes to the self the visual or auditory feedback of an action that was not performed by oneself (Banakou & Slater, 2014; Kokkinara et al., 2016; Wegner et al., 2004). Such illusory experiences of agency have been reported for example when participants feel high levels of body ownership over a surrogate body (Banakou & Slater, 2014; Kokkinara et al., 2016). Body ownership over virtual bodies can be induced by visuotactile (Slater et al., 2008) and visuomotor (Sanchez-Vives et al., 2010) correlations. Furthermore, ownership over a virtual arm can be induced through an SMR-based BCI, such that imagining a movement and seeing a virtual arm move from a first-person perspective is sufficient to induce certain levels of body ownership (Perez-Marcos, Slater, & Sanchez-Vives, 2009). Our findings are in line with the existing literature and expand the current knowledge on this topic: participants reported the highest feelings of body ownership when they induced the virtual movement through motor imagery; when inducing it through visually evoked potentials or when simply observing, the levels of body ownership were lower (see Fig. 2). Since in the present study self-attribution of the virtual body’s movement was mainly based on visual feedback, the role of body ownership is crucial for attributing an action to the self (Banakou & Slater, 2014; Kilteni & Ehrsson, 2017; Kokkinara et al., 2016; Wegner et al., 2004). Motor imagery and motor execution of the same movement are believed to activate similar neural networks (Ehrsson, Kuhtz-Buschbeck, & Forssberg, 2002; Gerardin et al., 2000; Pfurtscheller, Scherer, Muller-Putz, & Lopes da Silva, 2008). This could mean that the brain predicts not only the sensory feedback of a planned (and executed) action, but also the feedback of an imagined one. If this holds true, it could explain why all agency-related questionnaire items have the highest ratings when participants imagined the movement.

### The sense of responsibility for BCI actions

In order to feel responsible for the outcome of an action, the action should be performed by oneself. The sense of agency can give us this information and thereby creates the basis for a feeling of responsibility over that action (Haggard & Tsakiris, 2009). Experiencing high levels of agency over one’s own movements should therefore lead to high levels of responsibility. Based on this we measured in this study the sense of agency and the sense of responsibility, and expected both to show similar response patterns. Indeed, we found a similar pattern in agency and responsibility: for both, imagining the movement induces the highest levels, while observing the movement induces the lowest levels. Interestingly, however, we found a dissociation of agency and responsibility when the movement was induced by visually evoked potentials. While visually evoked potentials induced higher levels of agency compared with observation, and lower levels compared to imagining the movement, BCI through SSVEP did not induce higher responsibility levels than mere movement observation. Importantly, we also found that when imagining the movement, the level of activity in sensorimotor areas was positively correlated with the level of reported responsibility, meaning more sensorimotor activity was associated with higher responsibility ratings. It has been previously shown that when people see another person’s hand making movements at a position where they would expect their real hand to be, and when they are primed with auditory instructions given via headphones to make the hand movements, people feel illusory agency over the other person’s movements (Wegner et al., 2004). This priming can also be seen as an expectation of movement and when the participant creates an intention instead of the expectation, as is the case in the two BCI conditions, its effect should be at least similar, and possibly even stronger. Therefore, for responsibility, the combination of intention and seeing the virtual arm movement at an expected position is not sufficient. It has also been also suggested that for BCI actions the visual feedback dominates the sense of agency (Evans et al., 2015), meaning that poorly controlled BCI actions can result in a high sense of agency when the visual feedback matches the participant’s intended direction. In this experiment we manipulated the level of sensorimotor involvement and whether there is an intention or not. In line with Evans et al., (2015) our data shows that the sense of agency is higher when participants had formed an intention. However, our data further shows that when sensorimotor areas are involved in a BCI action participants report higher levels of agency and only then report responsibility. Therefore, in order to induce high feelings of responsibility it seems reasonable to suggest that the brain should create a motor plan.

### Therapeutic applications

The combination of virtual embodiment with motor imagery is also a promising approach for therapeutic applications. We now briefly discuss our results on this context. Observing, imagining, and actually performing the same motor action produces similar brain activity patterns, although with some differences (Jeannerod, 2001). Action observation and imagination are known to play an important role in motor learning (Feltz & Landers, 1983; Meltzoff & Moore, 1977). Therefore, action observation and imagination have been proposed as interventions for neurological rehabilitation (Buccino, Solodkin, & Small, 2006; Dickstein & Deutsch, 2007). Instead of promoting *compensation*, which bypasses the affected brain areas in order to achieve the intended action, action observation and imagination may instead induce recovery of motor deficits (Buccino et al., 2006), at least to some degree; this process is also called *remediation* (see Buccino (2014); Buccino et al. (2006), for reviews). *Remediation* processes are believed to add to existing therapy approaches. For instance, a study with stroke patients demonstrated that action observation combined with traditional physiotherapy improves motor function to a greater extent than in a control group that received only physiotherapy (Ertelt et al., 2007). Those patients reorganized their motor areas including the recruitment of a frontoparietal network similar to the action observation network, which might have given them a rehabilitation advantage (Ertelt et al., 2007). Motor imagery and body ownership as well as motor imagery, motor observation, and motor execution have been shown to have overlapping brain mechanisms (Evans & Blanke, 2013; Gallese, Fadiga, Fogassi, & Rizzolatti, 1996; Iacoboni et al., 1999), so they all activate sensorimotor areas through different networks. A combination of body ownership, motor imagery, and motor observation leads to higher experiences of agency and responsibility as shown in this study. It is still not clear, however, whether a combination of these three concepts, all of which have the ability to activate sensorimotor areas, would also foster *remediation* processes.

Simple motor imagery programs using, for example, hand laterality recognition tasks, imagined hand movements, and mirror therapy have been shown to be effective for complex regional pain syndrome (CRPS) (Llobera et al., 2013; Moseley, 2004). Motor imagery treatments have been suggested to reconcile motor output and sensory input, implying that any mismatch between motor intent and sensory feedback might be responsible for the pain felt in these conditions. Future work should focus on whether inducing higher levels of body ownership, agency, and responsibility by using a BCI to induce movements through activation of sensorimotor areas has a greater therapeutic pain-relieving effect than simple motor imagery programs in patients with CRPS and phantom limb pain.

BCI has been used in its different paradigms, and in particular motor imagery-based BCI, for the control of prosthetic limbs (Christoph Guger, Harkam, Hertnaes, & Pfurtscheller, 1999) and androids (Alimardani, Nishio, & Ishiguro, 2015). The sense of ownership, sensory feedback, and sense of control are critical for the successful integration of prostheses in the body representation.

### Ethical aspects

Remotely BCI-controlled androids are a promising possibility for the world and for the social interaction of immobilized patients, and the sense of control and responsibility over, for example, limb prostheses and more remote effectors (Tidoni, Gergondet, Kheddar, & Aglioti, 2014), should be a central consideration when developing these systems. This is a topic of high ethical relevance (Yuste et al., 2017), since we should expect in the next few decades a considerable growth of different forms of BCIs that can control anything from a screen cursor to a robot in another continent (Tidoni et al., 2014). A thorough understanding of the responsibility felt for these actions and its neural basis is therefore of the utmost importance.

## Data statement

Data is available from the corresponding author on reasonable request.

## Author contributions

MVSV, MM, and BN designed the experiment. MVSV supervised the different aspects of the study. BN and BS worked on the implementation and integration of the virtual reality and BCI. BN, MM and MB ran the experiments. BN, VVN and MVSV analyzed the data. BN and MVSV wrote the paper and all authors contributed to the final version.

## Acknowledgments

We would like to thank gtec Inc. for their support with the BCI integration, and Cristina Gonzalez-Liencres and Tony Donegan for editing.

