## Supplementary Material for "Agency and responsibility over virtual movements controlled through different paradigms of brain–computer interface"

### Supplementary Information

#### Supplementary Information Text

##### Methods

**Screening procedure.** A known limitation in SMR-based BCI protocols is that not all people can learn to control the BCI, especially not in less than 1 h (which was necessary to participate in the experiment). This issue is called BCI illiteracy and concerns 15-30% of participants (1). We encountered this problem in our pilots and therefore decided to pre-select our participants based on the clear occurrence of the sensorimotor alpha rhythm. Power spectral density in Laplace-transformed EEG-channels (C3 for right hand motor imagery) has been identified as predictor of SMR-based BCI performance (1). Therefore, we decided to pre-select participants based on the occurrence of a clear sensorimotor alpha rhythm. If we did not see a clear sensorimotor alpha peak in the participant's power spectrum, or if the first two CSP patterns, generated by our SMR-based BCI system, did not reflect activity in right hand and left foot sensorimotor areas, we excluded them from further participation in the experiment.

**Balancing.** The order of conditions was originally balanced over participants. Table S1 displays the frequency each condition was presented either as the first, second, or third condition.

**Table S1.** Frequency of one condition being presented as the first, the second, or the third condition.

| | First condition | Second condition | Third condition |
|---|---|---|---|
| ***Sample of 29 participants*** | | | |
| Observe | 9 | 10 | 10 |
| SSVEP | 12 | 8 | 8 |
| MotorImagery | 8 | 11 | 11 |
| ***Sample of 18 participants*** | | | |
| Observe | 6 | 4 | 4 |
| SSVEP | 7 | 6 | 6 |
| MotorImagery | 5 | 8 | 8 |

#### Results

**Questionnaire responses**. In addition to the principle components, we report in Table S2 Median values and inter-quartile ranges (IQR) for each condition.

**Table S2.** Median and IQR values of each questionnaire item in the different conditions (N = 29).

| Item tag | Observe | | SSVEP | | MotorImagery | | Phase |
|---|---|---|---|---|---|---|---|
| | Median | IQR | Median | IQR | Median | IQR | |
| *MyBody* | *10* | *1* | *10* | *1* | *10* | *1* | *Embodiment (Pre-condition)* |
| *IControlledArm* | *8* | *2* | *9* | *2* | *9* | *2* | *Embodiment (Pre-condition)* |
| *IControlledArm* | *5* | *6* | *8* | *4* | *9* | *3* | *Action* |
| *IBrokeObject* | *5* | *5* | *6* | *4* | *8* | *4* | *Action* |
| *Catch trial* | *2* | *4* | *2* | *3* | *4* | *6* | *Action* |
| *MyMovement* | *6* | *6* | *7* | *4* | *9* | *2* | *Post-condition* |
| *ArmMovedItself* | *4* | *6* | *8* | *5* | *9* | *1* | *Post-condition* |
| *MySound* | *3* | *5* | *7* | *3* | *8* | *4* | *Post-condition* |
| *ITouchedObject* | *5* | *4* | *6* | *5* | *8* | *5* | *Post-condition* |
| *MyBody* | *6* | *6* | *7* | *3* | *8* | *3* | *Post-condition* |

**IBrokeObject.** In each condition there were four trials in which an object fell off the table and broke. These trials were followed by the questionnaire item IBrokeObject. In three of these trials the virtual arm touched the object and caused it to fall off the table (Figure S4 A and B), while in one of these four trials the object fell by itself off the table without the virtual arm touching it (see Figure S4 C). This trial, which we refer to as the NoTouch trial, was used to check whether the IBrokeObject item was actually measuring responsibility. Ratings were significantly lower during the NoTouch trial. This was confirmed with a multilevel mixed-effects ordered logistic regression with fixed factor "trial" and "EMp" as covariate, which revealed a significant effect of "question" ($z = 7.12$, $P < .001$). Post-hoc paired comparisons showed that in the NoTouch-trial ratings were significantly lower than in the first (*Scheffé* $z = 6.28$, $P < .001$), the second (*Scheffé* $z = 6.41$, $P < .001$), and the third IBrokeObject trials (*Scheffé* $z = 7.12$, $P < .001$). There was no difference between second and first (*Scheffé* $z = 0.10$, n.s), third and first (*Scheffé* $z = 0.91$, n.s), or third and second IBrokeObject trials (*Scheffé* $z = 0.82$, n.s). Note that for this analysis the IBrokeObject ratings were downscaled to an ordinal scale ranging from 1 to 10.

**Embodiment phase.** The embodiment phase served to boost the feeling of body ownership over the virtual body. Participants reported high MyBody and IControlledArm ratings at the end of this phase, indicating that it induced high feelings of body ownership and control over the virtual body as intended by our experimental design. Neither IControlledArm ratings (ordered logistic regression, $z$ = 1.01, $P>0.05$) nor MyBody ratings ($z$ = -0.41, $P>0.05$) differed between experimental conditions during the embodiment phase.

**Covariates.** To control for possible misattribution of actually performed movements or muscle activity to the virtual movement, we introduced the following covariates: muscle activity (MA) in the right shoulder and eye movements (EM). We measured them during the preparation phase (MAp and EMp) and during the motor observation phase (MAo and EMo) and added them as covariates to our statistical model in all agency-related measures. We kept them in the model if they improved the fit of the model to the data. MA and EM differed in the three experimental conditions in the preparation phase (MAp: $F(2,52)$ = 3.81, $P$ = .029, EMp: $F(2,56)$ = 3.54, $P$ = .036), and in the motor observation phase (MAo: $F(2,52)$ = 2.78, $P$ = . 071, EMo: $F(2,56)$ = 4.59, $P$ = .014).

**Event-related synchronization.** We calculated ERS in occipito-parietal areas during the SSVEP condition in the same way as ERD with just two changes: We filtered our data between 6-10 Hz, a range that includes both SSVEP frequencies, and we selected the strongest CSP-patterns for ERS in visuo-parietal areas. There was no relation between neither ERS% and agencyPCF (z = 0.37, P = 0.715) nor ERS% and responsibilityPCF (z = 0.41, P = 0.682) as tested with a mixed-effects regression.

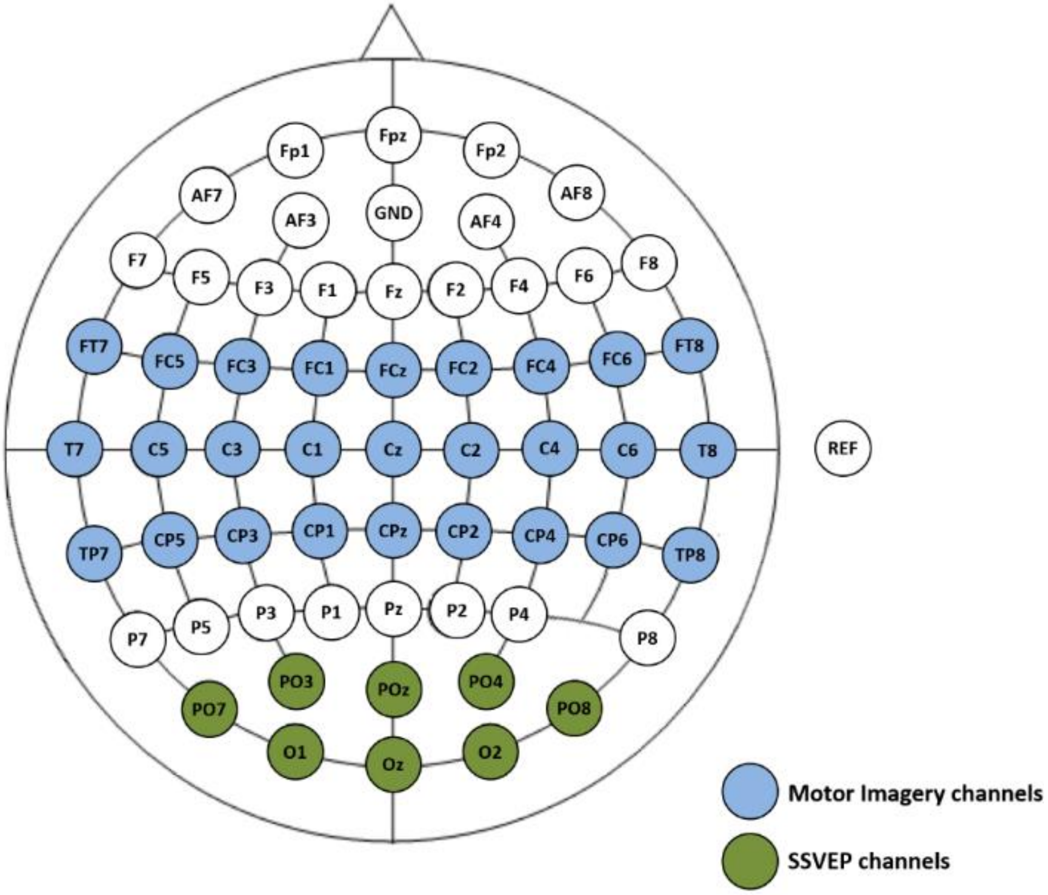


**Fig. S1.** EEG montage. The electrode positions used in this experiment are displayed. Ground was positioned at AFz and electrodes were referenced to the right ear lobe. Colored electrodes indicate electrodes used for online analysis—blue for SMR-based BCI and green for SSVEP-based BCI.

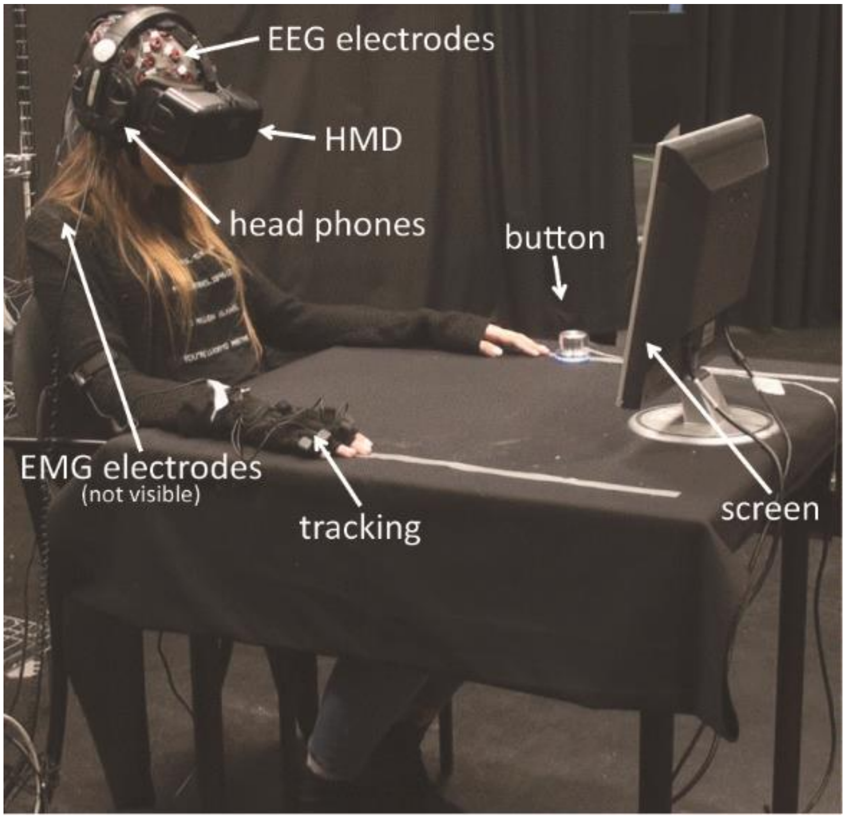


**Fig. S2.** Experimental setup. Position of the participant throughout the experiment. Participants were asked to move their right hand during the embodiment phase and movements were tracked and mapped onto the virtual right arm. During the other parts of the experiment, they were asked not to move their hands. Instructions were given via headphones.

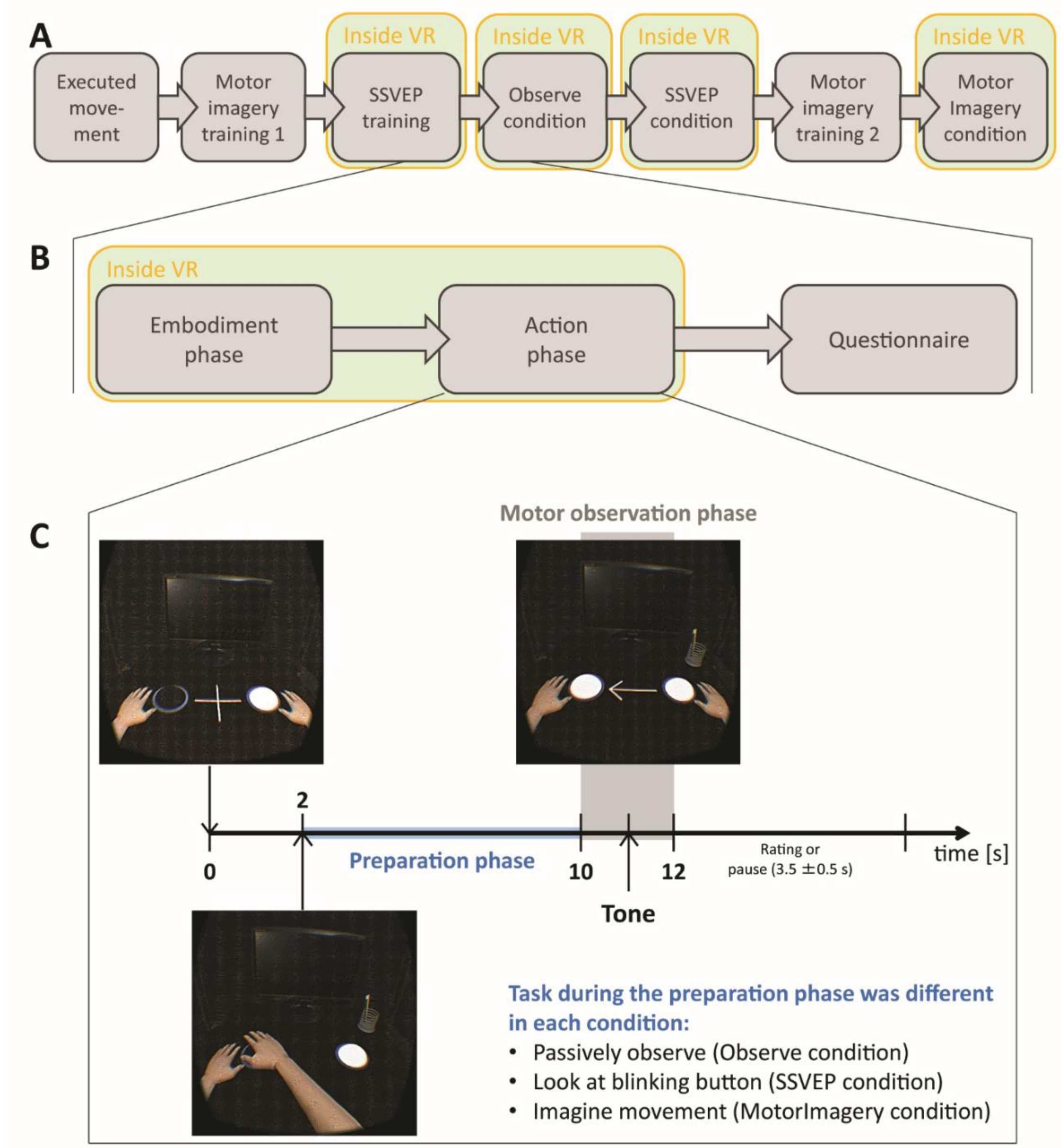


**Fig. S3.** Timeline of the experimental procedure. (A) Timeline of the whole experiment. Motor imagery training was divided into two parts—one at the beginning and one just before the MotorImagery condition started. The order of conditions was randomized among participants. (B) Timeline of one experimental condition. Each condition started with the embodiment phase followed by the action phase, both of which were inside virtual reality (VR), and ended with the completion of a questionnaire. (C) Timeline of one trial. Each experimental trial started with a fixation cross followed by an arrow indicating the button to which the arm should move. This was followed by a preparation phase in which participants either passively observed what was happening (Observe), looked at the indicated blinking button (SSVEP) or imagined the arm movement to this button (MotorImagery). This was followed by the movement of the arm (or an error tone if the BCI classifier did not detect the movement). Every 10 trials participants had to rate their experience of control over the virtual movement (item IControlledArm).

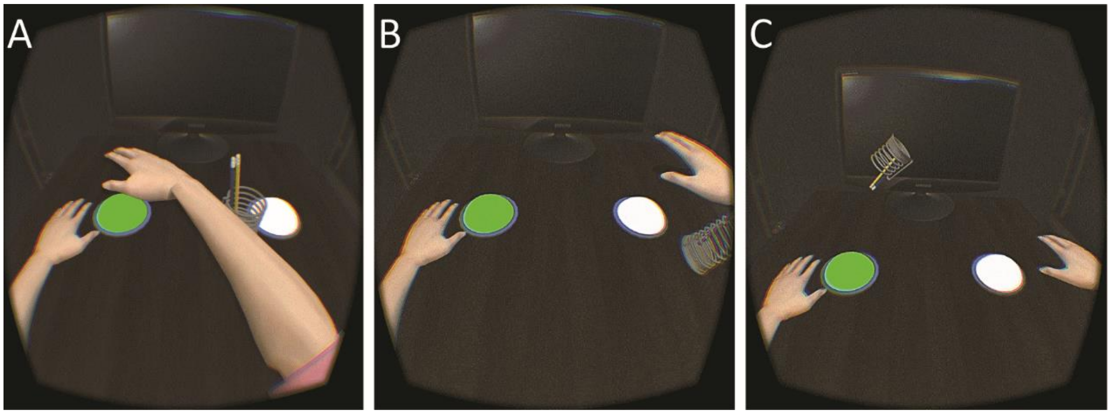


**Fig. S4.** Responsibility trial. In four random trials during each experimental condition, an object fell off the table. In three of those trials the virtual body threw the object off the table (A and B) and in one trial the object fell of the table by itself (C). The object was different in each of the four trials, alternating between a pencil stand, a vase, a coffee cup and a glass. Participants were asked to rate the degree of perceived responsibility (item IBrokeObject) directly after these trials.

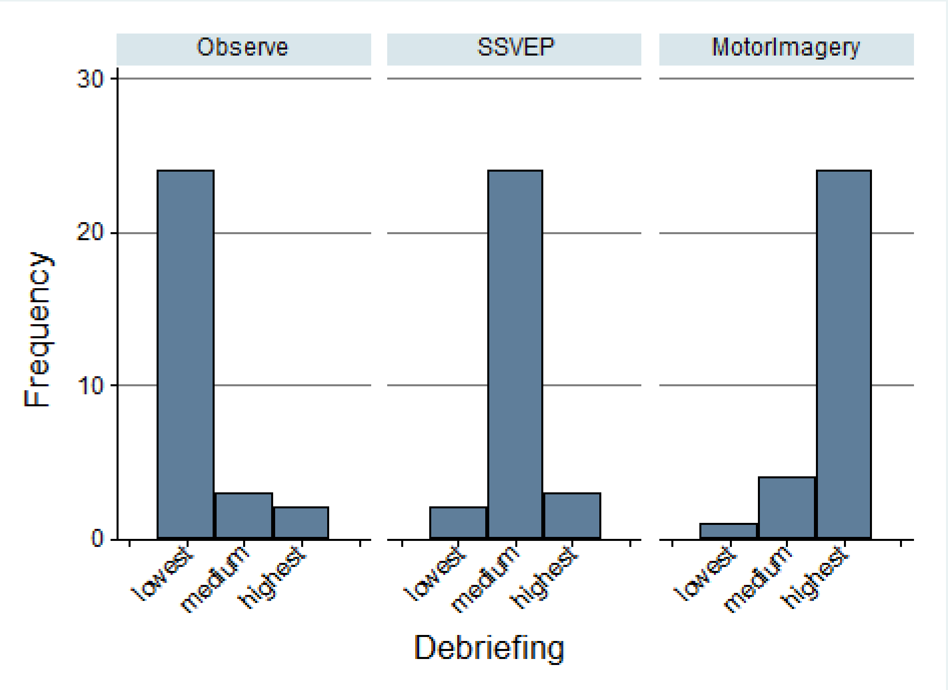


**Fig. S5.** Histogram of debriefing responses. Participants were asked to order the conditions regarding the sense of control they perceived during the condition over the virtual arm movement from high to low.
